# Multiband entropy-based feature-extraction method for automatic identification of epileptic focus based on high-frequency components in interictal iEEG

**DOI:** 10.1101/2020.03.23.004572

**Authors:** Most. Sheuli Akter, Md. Rabiul Islam, Yasushi Iimura, Hidenori Sugano, Kosuke Fukumori, Duo Wang, Toshihisa Tanaka, Andrzej Cichocki

## Abstract

Presurgical investigations for categorizing focal patterns are crucial, leading to localization and surgical removal of the epileptic focus. This paper presents a machine learning approach using information theoretic features extracted from high-frequency subbands to detect the epileptic focus from interictal intracranial electroencephalogram (iEEG). It is known that high-frequency subbands (>80 Hz) include important biomarkers such as high-frequency oscillations (HFOs) for identifying epileptic focus commonly referred to as the seizure onset zone (SOZ). In this analysis, the multi-channel interictal iEEG signals were splitted into segments and each segment was decomposed into multiple high-frequency subbands. The different types of entropy were calculated for each of the subbands and the sparse linear discriminant analysis (sLDA) was applied to select the prominent entropy features. Due to the imbalance of SOZ and non-SOZ channels in iEEG data, the use of machine learning techniques is always tricky. To deal with the imbalanced learning problem, an adaptive synthetic oversampling approach (ADASYN) with radial basis function kernel-based SVM was used to detect the focal segments. Finally, the epileptic focus was identified based on detection of focal segments on SOZ and non-SOZ channels. Eight patients were examined to observe the efficiency of the automatic detector. The experimental results and statistical tests indicate that the proposed automatic detector can identify the epileptic focus accurately and efficiently.

## Introduction

Epilepsy is one of the most common neurological disorders of the nervous system, affecting people worldwide at any age. According to the World Health Organization (WHO), approximately 50 million people globally have been diagnosed with epilepsy, which causes social impairment and carries a higher risk of death ^1, 2^. Epilepsy is defined as repeated and unpredictable seizures caused by abnormal neuronal firing in the brain ^3^. Physicians distinguish the type of seizures as either focal (partial) or generalized, based on the location of abnormal brain activity and its propagation ^4, 5^. Most patients are prescribed inexpensive daily medication to control epileptic seizures, but some become resistant to them; thus, resectioning of the epileptic focus surgically may provide the best chance of seizure control ^6, 7^. Therefore, the localization of the epileptic focus is crucial for epilepsy treatment. The standard diagnostic modalities for epileptic focus detection are the investigation of seizure semiology, MRI, and EEG. When the epileptologist cannot determine an epileptic focus after using noninvasive methods, the implantation of intracranial electrodes to record iEEG during both of the interictal and ictal phases is indicated.

In practice, accurate detection of the epileptic focus is generally achieved by epileptologists observing long-term iEEG categorizing the patterns of the seizures. The visual examination of long-term iEEG is a time-consuming and laborious process, as the detection of the seizures from the interictal time in iEEG is the most difficult task ^8, 9^, which puts a heavy burden on epileptologists and reduces their efficiency. Therefore, a computer-aided system with an effective algorithm that uses the iEEG signal to localize the epileptic focus would be invaluable.

Recent studies have proposed various machine learning-based methods for the problem of identifying focal and nonfocal iEEG signals ^10–13^. Typical methods extract appropriate features that reflect the dynamics that characterize normal and epileptic brain activities. Feature extraction is followed by a classifier such as linear discriminant analysis (LDA), k-nearest neighbor (kNN), or SVM. For machine learning-based methods, information theoretic features, particularly entropies, have been established as efficient features. Srinivasan et al. proposed the use of approximate entropy (AE) as a feature of artificial neural networks (ANNs) for automatic seizure prediction ^14^. Nicolaou et al. ^15^ proposed a permutation entropy (PE) to classify epileptic signals from iEEG signals to use with SVM. Song et al. used sample entropy as a feature extraction method for detecting healthy vs epileptic seizures ^16^. Recently, a variant of time domain multiband decomposition methods, which include empirical mode decomposition (EMD) and bivariate EMD, has been proposed to improve the accuracy of seizure detection in combination with the different types of entropy-based feature-extraction approaches ^17, 18^. The studies showed evidence that supports the use of entropy features to characterize normal and epileptic activities. Although the above methods have achieved promising results in detecting the epileptic events, these studies are restricted by the limited pairs of channels as well as lower frequency bands (0.5–150 Hz) used Bern-Barcelona and other datasets ^10, 19–21^.

However, the use of a low frequency band that partially excludes higher frequency components has some restrictions in real world applications. Recent studies in epilepsy ^22,23^ have shown that high-frequency oscillations (HFOs), including ripple (80–250 Hz) and fast ripple (250–600 Hz), are promising biomarkers of the epileptic focus detection. It has been shown that fast ripple consists of normal brain activities associated with visual perception, which also complicates the clinical use of HFOs as valid biomarkers to guide epilepsy surgery ^22–24^.

In some EEG-based studies ^25–28^, due to the non-linear and non-stationary properties of signals, the filter-bank approaches were introduced to improve the performance. Several machine-learning and statistical methods were combined with the multiband framework to extract noise-robust features from the narrow band signals ^25–27^. In this study, we hypothesize that this filter-based approach is effective for identifying epileptic focus using a high frequency bands. The aim of this study is to identify epileptic focus (more specifically Seizure Onset Zone; SOZ) ^102^ using multi-band entropy-based features with machine learning in high frequency components of interictal clinical iEEG. To the best of our knowledge, multi-band entropy-based feature extraction in conjunction with high-frequency components (ripple and fast ripple) has not been reported for use in the detection of epileptic focus from continuous interictal iEEG signals. Considering the noise-robust features and the reduction of system complexity, the multi-band feature-extraction method has a great potential as the basis for designing a computer-aided system for localizing epileptic focus. The rest of this paper is organized as follows: Result section presents the simulated experiments with the proposed methods using iEEG data from eight epilepsy patients. The detailed description of the proposed automatic system used in the clinical situation, comparison with existing method to evaluate the proposed system, and future works are presented in the Discussion section. The conclusion section presents the conclusion of this manuscript. The materials and methods section is presented the dataset, data pre-processing, multiband analysis, entropy feature-extraction, feature selection, imbalanced learning problem, cross-validation design, and performance measurements metrics.

## Results

First, the performances based on different algorithms, including filter-bank approach (FbA), FbA with ADASYN (FbA/ADA), and FbA with feature selection and ADASYN (FbA/FS/ADA), were compared with 10-fold cross-validation. Second, the performances of different approaches were considered and the optimum one was selected to simulate the results for eight epilepsy patients. The parameters for the proposed algorithms were summarized as follows:

**Algorithm 1-(FbA):** In this case, the *N* bandpass filters were implemented using a third-order Butterworth filter to subband the high-frequency components in iEEG. The different types of entropy measuring methods were applied onto each subband to extract features. The extracted features were input to SVM with a 10-fold cross-validation for classifying the focal and non-focal segments. Of note, the feature selection and oversampling technique (ADASYN) were not used.
**Algorithm 2-(FbA/ADA):** The *N* bandpass filters and feature extraction were performed in the same way as in **Algorithm 1**. For handling the imbalanced learning problem, the ADASYN method was used in the training stage of the SVM for each cross-validation.
**Algorithm 3-(FbA/FS/ADA):** In this algorithm, subbanding and feature extraction were implemented in the same way as in **Algorithm 1**. The selection of dominant entropy features was performed using sparse LDA. Finally, the selected features were input to the SVM for classifying focal segments.

### Effect of Feature Selection

The combination of eight entropies were used to extract features from each subband defined in eq. (16). From the eight entropy features, we hypothesized that some features would be more effective for the purpose of recognition. Therefore, we used sLDA weights from the training set induced in eq. (17) to select the prominent features based on non-zero weights from each subband. In this study, we set the sparsity parameters *δ* = 3 and *δ*_1_ (Pt1: *δ*_1_ = −5; Pt2: *δ*_1_ = −5; Pt3: *δ*_1_ = −3; Pt4: *δ*_1_ = −5; Pt5: *δ*_1_ = −3; Pt6: *δ*_1_ = −4; Pt7: *δ*_1_ = −6; Pt8: *δ*_1_ = −3) based on the training set to achieve satisfactory results, where the absolute value of *δ*_1_ corresponds to the desired number of variables. Fig. 1 shows the colormap of sLDA weights as a function of subbands on the vertical-axis and the different types of entropies on the horizontal-axis. The figure indicates that the features with higher values of weights are more significant. To justify this hypothesis, the average area under the ROC curve (AUC) was derived with 10-fold cross-validation from individual entropy features, as well as the average weights across *N* subbands, were estimated, as illustrated in Fig. 2. This figure shows the relationship between the weights of entropies and the AUC of individual entropies suggesting that the entropies with non-zero weights may improve the system performance. Therefore, to select the prominent features for the purpose of classification, the entropies corresponding to non-zero weights were selected to evaluate the performance of the system.

**Fig. 1.**
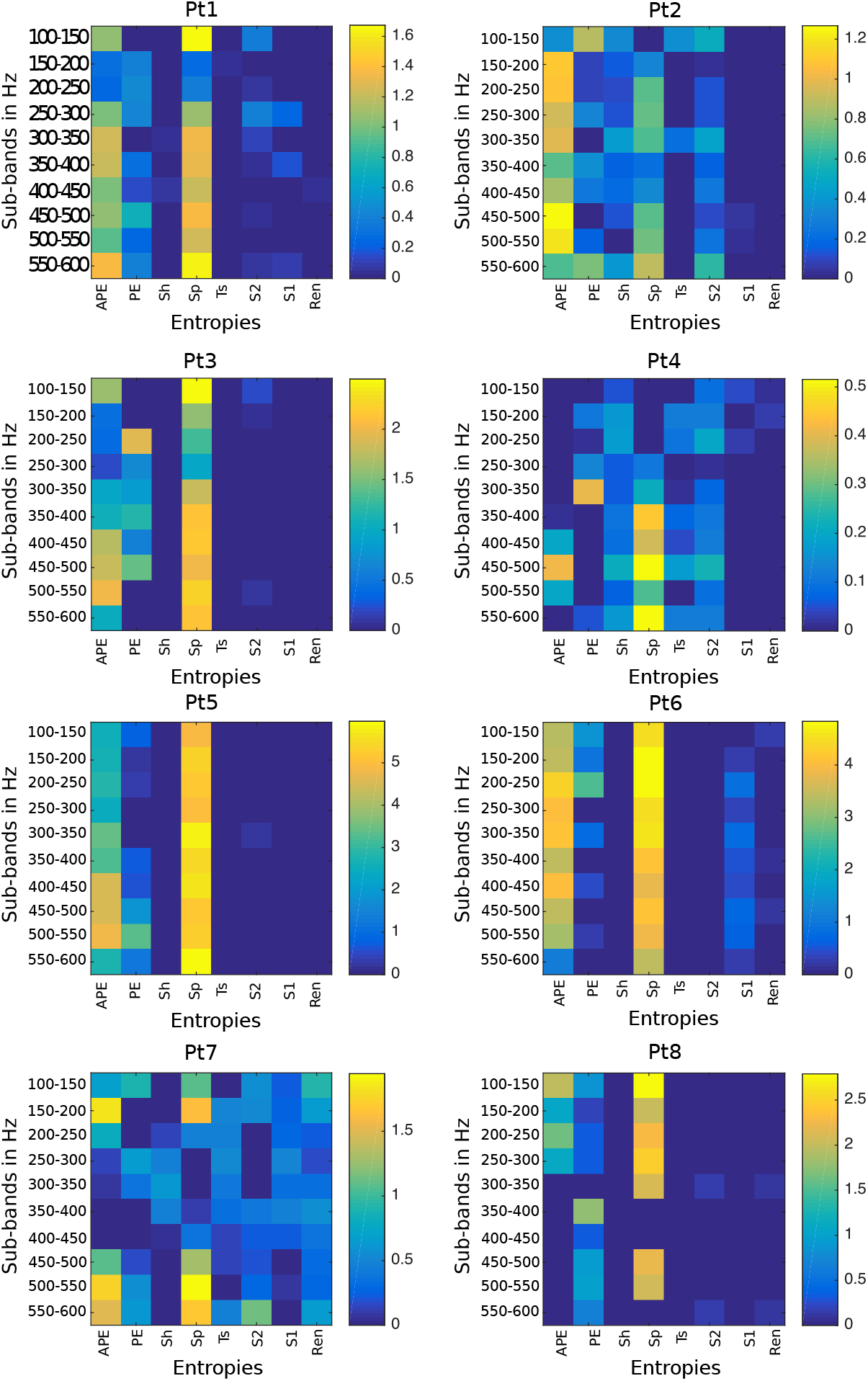
The color map representing the sLDA weights of the entropies with each subband for eight patients. The entropies represented in this paper are: APE (Approximate Entropy), PE (Permutation Entropy), Sh (Shannon Entropy), Sp (Sample Entropy), Ts (Tsallis Entropy), S2 (Phase Entropy 2), S1 (Phase Entropy 1), and Ren (Reny’s Entropy).

**Fig. 2.**
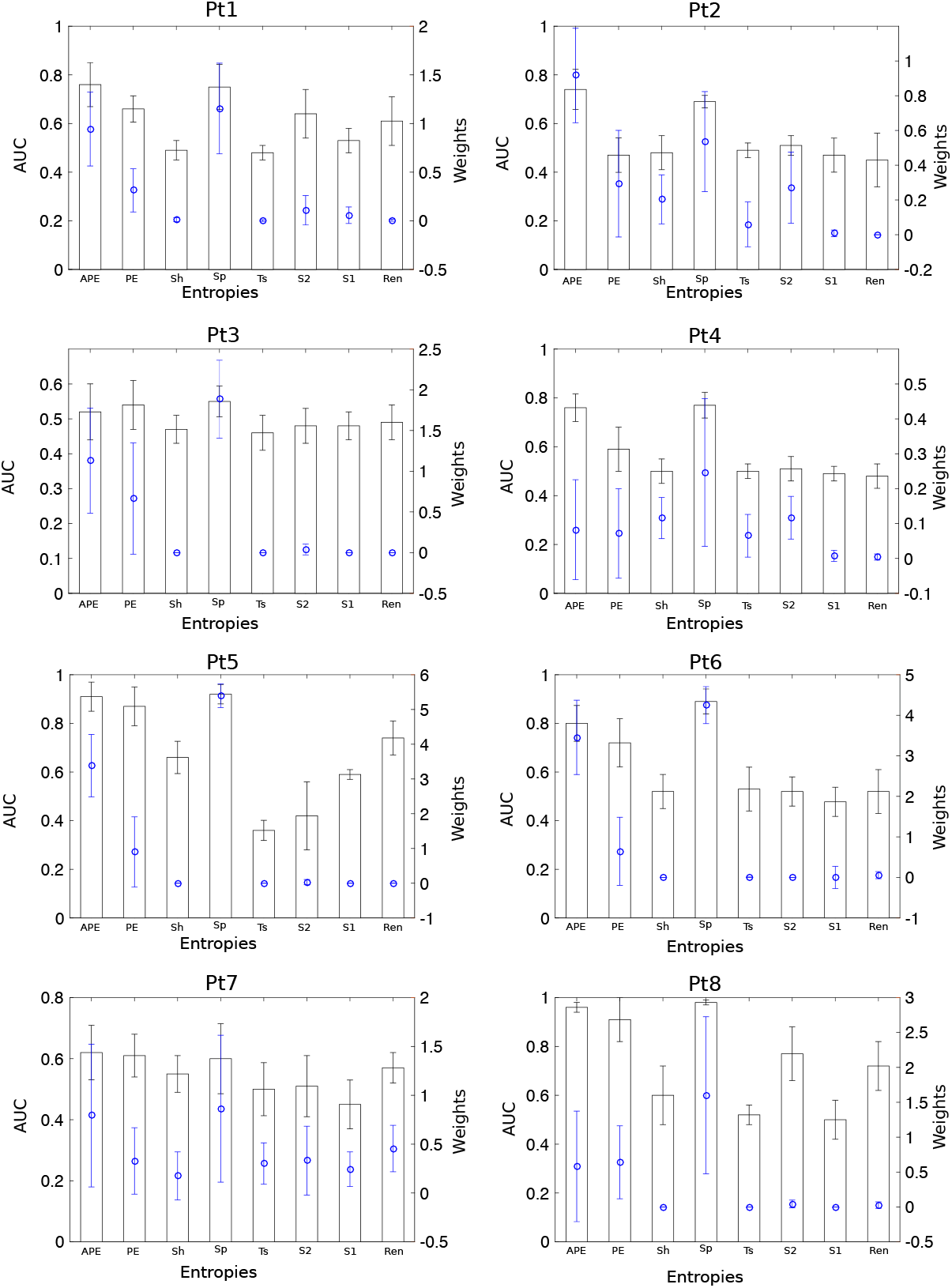
Average AUC obtained from an individual entropy feature (bar) and its average sLDA weights across subbands (blue). Error bars indicate standard errors.

### Performance Analysis with Different Cases

To evaluate the performance in the different algorithms (FbA, FbA/ADA, and FbA/FS/ADA), the AUC was performed for eight patients shown in Table 1. The AUC of a method in cases of imbalanced learning is equivalent to the probability of ranking a randomly chosen positive instance higher than a randomly chosen negative instance ^29^. In this experiment, we set all parameter for ADASYN according to the study ^30^ to balance the training features. The AUC for the algorithm FbA/ADA with feature selection exhibits superior results for all eight patients. The reason for lower performance using the FbA method is that the high degree of imbalance distribution between the minority (focal segments) and majority (non-focal segments) class may provide biased decision boundary used in SVM training. In the test of the statistical significance of the methods, the result of Friedman’s ANOVA showed a significant main effect on AUC (*p* < 0.05). Performing a Tukey-Kramer-based post-hoc test, the method using FbA/FS/ADA achieved significantly higher AUC across all eight patients than the other methods (FbA vs FbA/ADA: *p* < 0.001; FbA/ADA vs FbA/FS/ADA: *p* < 0.001).

**Table 1:** Area Under the ROC Curve (AUC) comparison with different cases (FbA, FbA/ADA, and FbA/FS/ADA) for eight patients. The average AUC is estimated for individual segments with 10-fold cross-validation.

| Patient ID | FbA | FbA/ADA | **FbA/FS/ADA** |
| --- | --- | --- | --- |
| Pt1 | 0.64 | 0.77 | **0.79** |
| Pt2 | 0.71 | 0.78 | **0.83** |
| Pt3 | 0.52 | 0.52 | **0.63** |
| Pt4 | 0.66 | 0.73 | **0.75** |
| Pt5 | 0.93 | 0.95 | **0.97** |
| Pt6 | 0.90 | 0.94 | **0.97** |
| Pt7 | 0.65 | 0.70 | **0.70** |
| Pt8 | 0.96 | 0.98 | **0.99** |
| Mean | 0.74 | 0.79 | **0.83** |

### Results with focal Segments-spotting

In this section, we provide the results of individual focal segment detection based on the optimal algorithm (FbA/FS/ADA) with eight epilepsy patients. The result showed in the above section that the selection of prominent features and the oversampling method can significantly improve the performance of an automatic system. Hence, we are the first one to use high-frequency components (ripple and fast ripple) from interictal iEEG, the performances of localizing individual segments were observed in terms of sensitivity, specificity, precision, fall-out, and F-score for the comparison study similar to HFOs- and low frequency-based related studies ^10, 19–21, 31–35^ computed from the confusion matrix (shown in Table 6). It is observed from the Table 2 that the proposed method achieved the highest performance for localizing individual segments with the adult patients Pt5 (SEN: 79.25%; fall-out: 2.50%), Pt6 (SEN: 54.82%; fall-out: 3.58%), and Pt8 (SEN: 88.52%; fall-out: 1.46%). The positive likelihood ratios (PLRs) were also used to evaluate the studies in different research ^36–38^. The PLRs for the adult patients (Pt5, Pt6, and Pt8) were achieved 31.70, 15.31, and 60.63, respectively. For the deep sheeted FCD patients (Pt2 and Pt7), the surgeon implanted the small electrodes vertically on the sulcus. The sensitivity and Fall-out were Pt2 (SEN: 45.93%; fall-out: 13.36%), Pt7 (SEN: 49.02%; fall-out: 16.24%) and the PLRs were achieved 3.44 and 3.02 for patients Pt2 and Pt7, respectively. In case of pediatric patients, the sensitivity and Fall-out were Pt1 (SEN: 23.70%; fall-out: 3.74%), Pt3 (SEN: 37.46%; fall-out: 17.18%), and Pt4 (SEN: 42.96%; fall-out: 15.93%). The proposed method provided the PLRs of 6.34, 2.18, and 2.70 for the patients Pt1, Pt3, and pt4, respectively.

**Table 2:** Experimental results for individual segments using the optimal method (FbA/FS/ADA).

| **Patient ID** | **SEN [%]** | **SPE [%]** | **Precision [%]** | **Fall-out [%]** | **F-score** |
| --- | --- | --- | --- | --- | --- |
| **Pt1** | 23.70 | 96.26 | 25.00 | 3.74 | 0.24 |
| **Pt2** | 45.93 | 86.64 | 43.01 | 13.36 | 0.44 |
| **Pt3** | 37.46 | 82.83 | 30.37 | 17.18 | 0.34 |
| **Pt4** | 42.96 | 84.07 | 19.69 | 15.93 | 0.27 |
| **Pt5** | 79.25 | 97.50 | 62.57 | 2.50 | 0.70 |
| **Pt6** | 54.82 | 96.42 | 58.96 | 3.58 | 0.57 |
| **Pt7** | 49.02 | 83.76 | 27.71 | 16.24 | 0.35 |
| **Pt8** | 88.52 | 98.54 | 71.34 | 1.46 | 0.79 |
| **Mean** | **52.70** | **90.75** | **42.33** | **9.24** | **0.46** |

### Results with Channel Identification

Fig. 3 shows a graphical representation with the localization of the focal and non-focal segments, which help the epileptologists in two ways: (1) to observe the localization of the focal and non-focal segments over duration of the iEEG, and (2) the number of detected focal segments corresponding to the seizure onset and non-seizure onset channels. This provides useful information about the active electrodes located close to the epileptic focus. The vertical axis in the color map (left) represents the electrodes and the horizontal-axis shows the segment index. Each yellow spot in the color map represents the detected focal segments. The right side of the color map in each patient indicates the number of detected focal segments (*x*-axis) in each electrode (*y*-axis) in which a group of bars (red) represents “SOZ” and black bars without color indicates the “non-SOZ.” It is observed from the Fig. 3 that a sharp yellow spotted areas are clearly visible for each electrode of SOZ for the adult patients Pt5, Pt6, and Pt8. The detected focal segments (yellow spot) for the patients Pt2 and Pt7 with deep sheeted FCD were widely distributed through the non-SOZ. The possible reason of widely distribution was that the patients was the BOS-type FCD in which surgeon implanted vertical electrodes into the deep in the brain. A similar scenario was observed in the case of pediatric patients (Pt1, Pt3, and Pt4). Table 3 shows the results of identifying the epileptic focus for each patients by measuring the AUC across all possible thresholds based on detected focal segments in the SOZ and non-SOZ (see in Fig.3).

**Fig. 3.**
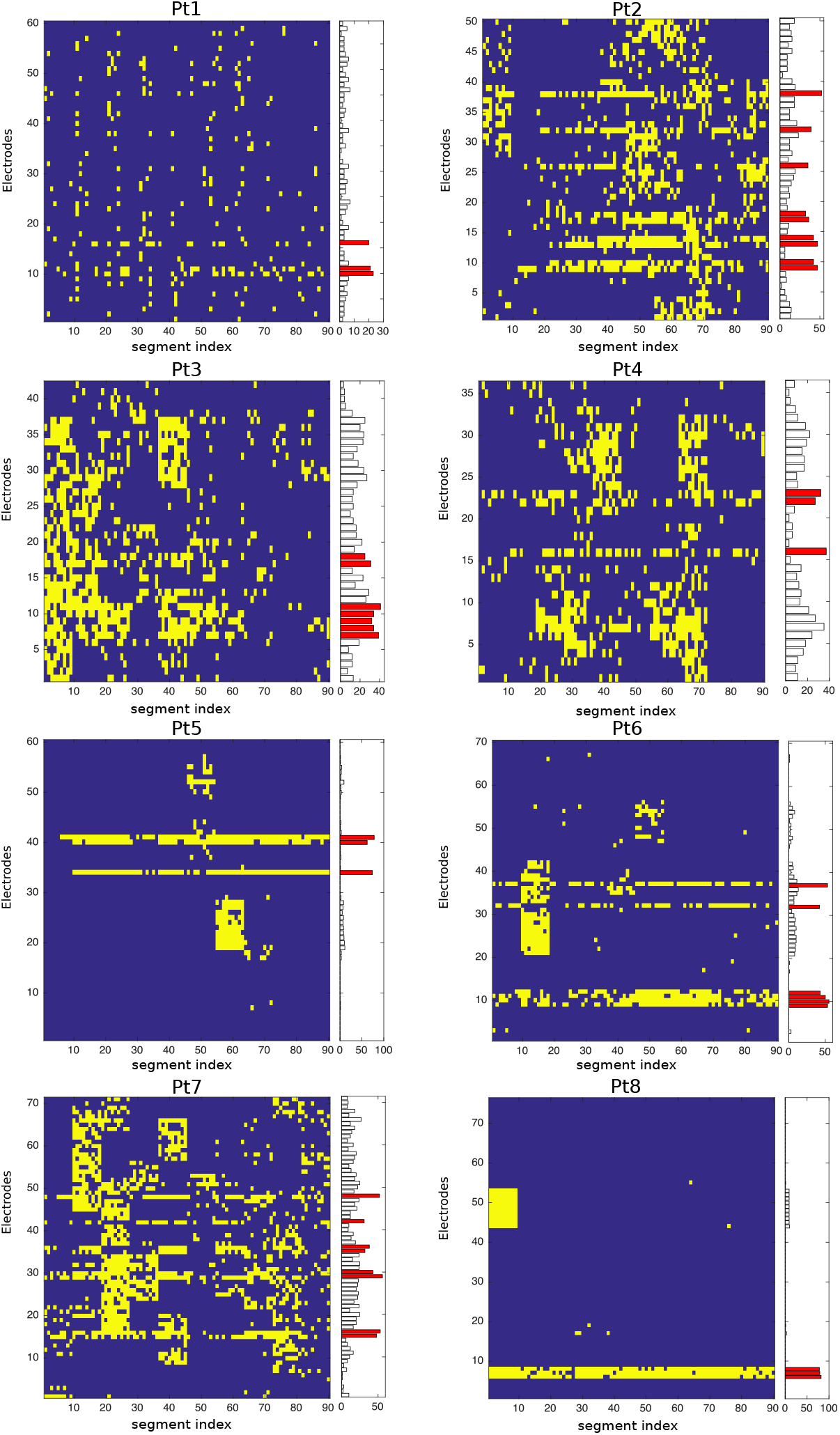
Color map representing the localization of segments (yellow spots) with respect to channels for the eight patients using our proposed method. The bar with each color map represents SOZ (red) and non-SOZ (black) with number of detected focal segments.

**Table 3:** Average AUC with 10-fold cross-validation for identifying epileptic focus.

| **Patient ID** | **Pt1** | **Pt2** | **Pt3** | **Pt4** | **Pt5** | **Pt6** | **Pt7** | **Pt8** | **mean** |
| --- | --- | --- | --- | --- | --- | --- | --- | --- | --- |
| **AUC** | 0.90 | 0.79 | 0.71 | 0.79 | 0.96 | 0.94 | 0.81 | 0.99 | **0.86** |

### Computational Cost

The average computational time for each entropy with 10 subbands was measured using Python on iMac Pro (with Intel Xeon W processor and 128 GB RAM). Note that the average results are estimated with 100 runs at the testing phase to detect a single segment. Table 4 shows the average computational time (in seconds) at each entropy with 10 subbands. It is observed that the phase entropy requires the highest computational time. The entropy with Ts, Ren, Sh, and PE requires shorter time compared to others. Since, we used the combination of all eight entropies to design the whole system, the average computational time with eight entropies and 10 subbands was 56.51 s to test a single segment.

**Table 4:**
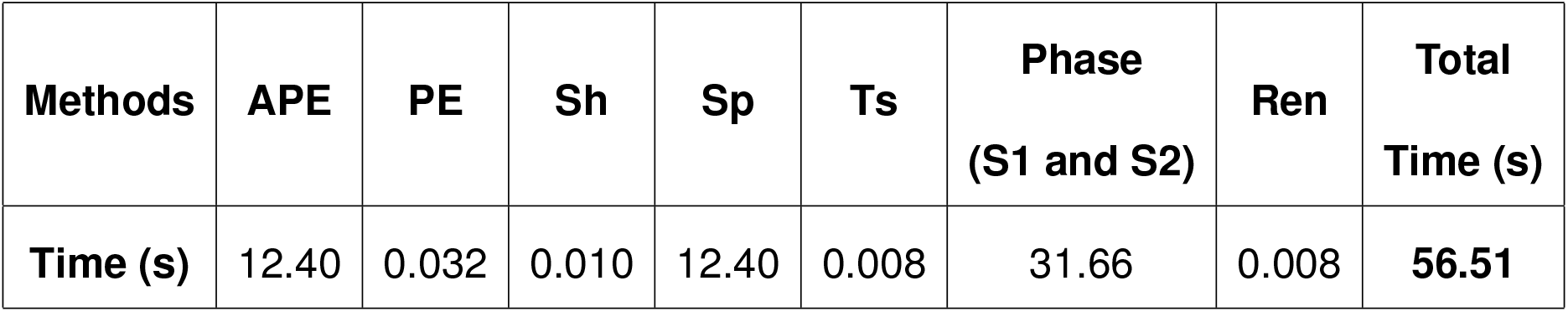
Average computational time (s) with each entropy for 10 subbands.

| **Methods** | **APE** | **PE** | **Sh** | **Sp** | **Ts** | **Phase (S1 and S2)** | **Ren** | **Total Time (s)** |
| --- | --- | --- | --- | --- | --- | --- | --- | --- |
| **Time (s)** | 12.40 | 0.032 | 0.010 | 12.40 | 0.008 | 31.66 | 0.008 | **56.51** |

## Discussion

According to the clinical guidelines related to epilepsy surgery, epilepsy surgeon should consider implanting the intracranial electrodes to observe the seizure onset zone (SOZ), irritative zone, and symptomatic zone before the epileptic focus resection for patients with medically intractable epilepsy. The epileptic zone includes SOZ and a part of irritable zone and symptomatic zone. To determine the epileptic focus, the epileptologists need to analyse and label 3 to 7 days iEEG data depending on the patients conditions. In the proposed automatic system, we need to label only 30-minutes of the interictal iEEG to localize the epileptic focus, instead of 3 to 7 days labeling. An automatic detection or estimation of SOZ from short period of interictal recording provide epileptologists a great assistance and can increase the number of iEEG analysis for patients with intractable epilepsy. A recent study developed EPINETLAB, a multi-graphic user interface (GUI) automated software, to help researchers and clinicians to detect HFOs and identify the SOZ using iEEG/MEG data ^50^. To perform a preliminary validation analysis of EEG data, they used six patients with drug-resistant epilepsy and analyzed only the ripple frequency band (80–250 Hz). However, a number of recent studies have found that fast ripple (200–600 Hz) could be more valid and reliable biomarkers than ripple bands to guide epilepsy surgery ^23, 24, 45^.

Several studies exploiting Bern-Barcelona and Bonn EEG datasets ^10,19^ have been reported for epilepsy-related signal classification. For instance, the Bonn datasets consist of five EEG datasets denoted as Set A (normal: healthy awake and eyes open), Set B (normal: healthy awake and eyes closed), Set C (Epileptic: interictal), Set D (Epileptic: interictal), and Set E (epileptic: ictal) with 100 single-channels and the time duration of each channel was 23.6 s. Nicolaou et al. used approximate entropy as a feature and employed SVM classifier for identifying normal vs ictal EEG with average accuracy of 93.55% ^15^. Guo et al. proposed an automatic epileptic seizure detection system with approximate entropy features derived from multi wavelet transform, and combined with an artificial neural network to classify the existence or absence of seizure with average accuracy 99.85% for two cases: (normal vs ictal), and the combination of normal and interictal vs ictal ^51^. In another study, wavelet packet entropy and hierarchical EEG classification were proposed with average accuracy of 99.44% for normal vs ictal ^52^. A method based on discrete wavelet transforms (DWT) with entropy features was proposed, leading to a classification accuracy of 84% using *k*-nearest neighbor (kNN), probabilistic neural network (PNN), fuzzy classifier, and least squares support vector machine (LS-SVM) ^53^. Mursalin et al. proposed an automated epileptic seizure detection approach using improved correlation-based feature selection and random forest classifier (RFC) with average accuracy of 98.44% ^54^. However, the studies with a variant of EEG and iEEG datasets including Friburg ^20^, CHB-MIT ^21^, Children’s hospital Boston datasets ^55^ etc. were investigated to identify the seizure events based on machine learning approaches. All the above studies used only lower frequency bands (0.5–150 Hz) for limited pairs of electrodes with well balanced problems. In this study, we have considered the full clinical perception, including high-frequency components (100–600 Hz) and multi-channels imbalanced problem, which offers the practical implementation of clinical utilization. Due to the highly imbalanced problem of the iEEG data, we used AUC here instead of using classification accuracy as system evaluation criterion. The proposed method detects epileptic focus for different epilepsy mechanisms patients (age and pathological type) with an average AUC of 0.86. We also observed that the entropy features, such as APE, PE, and Sp, are more discriminative in high-frequency bands. These findings may provide an excellent tool when appropriate methodology will be combined with the high- and low-frequency bands to locate the epileptic focus.

Recently, Ullah et al. has proposed an automated system for epilepsy detection for Bonn dataset based on deep learning approach, yielding 99.1% accuracy ^56^. A similar dataset was used to design a deep convolutional neural network (CNN) with 13 layer for categorizing the normal, preictal, and seizure class and obtained an average accuracy of 88.7%, a specificity of 90% and a sensitivity of 95% ^57^. Although, the deep-learning based automatic systems have improved the system performance compared to simpler classier SVM, it needs a large amount of training data in order to show such remarkable performance. On contrary to the deep learning, the SVM method is easy to understand and provides consistent performance. The epileptologists can efficiently interpret the classifier outcome to take the right medical decision.

A comparison study with Bern-Barcelona dataset using time-domain multiband analysis, including EMD and BEMD, was reported by Itakura et al. ^58^, improving the performance of the system with a 86.89% average accuracy for identifying the seizure patterns. However, this type of analysis is only suitable for single and bivariate iEEG signals. In the case of multi-channel iEEG signals, the number of extracted bands over channels are not consistent. Thus, the EMD and BEMD methods are not suitable for multi-channel iEEG signals. Considering real time implementation, a filter-bank technique is more convenient for decomposing high-frequency bands (100–600 Hz), has little computational cost, and decreases the system complexity when compared with other multiband approaches.

In epilepsy studies for identifying epileptic focus, the visual inspection of iEEG time series have demonstrated that HFO may occur during ictal, preictal, and interictal states ^39–42^, and the rate of HFOs tends to be higher in SOZ ^42–45^. To detect HFOs, several automatic HFO detectors have been proposed including the methods of artifact rejection, estimating the energy of the signal using Root Mean Square (RMS) amplitude, short-time Linelength or others ^46–49^. However, HFOs-related studies to identify the possible seizure onset channels need the long-time iEEG data to calculate the baseline. Jrad et al. proposed automatic HFO detection with multi-class SVM in depth-EEG signals ^59^. In their study, the performance evaluation matrices for evaluating the system were used in terms of sensitivity and false discovery rate (FDR). The reason for using FDR was that the amount of true negative (TN) was large enough in HFOs detection task. They achieved an average result with five drug-resistant epilepsy for Ripple (Sensitivity: 81.1% and FDR: 30.2%) and fast ripple (Sen: 74.6% and FDR: 6.3%). Guo et al. proposed magnetoencephalography-based (MEG) HFOs detector using stacked sparse autoencoder (SSAE) for identifying the HFOs and normal control (NC) with well balanced problem achieving 89.9% in accuracy^60^. The method CNN was used by Johansen et al. ^61^ for identifying spikes and HFOs with five epilepsy patients with an average AUC of 0.94. To detect spikes, ripples, ripples-on-spikes (RonS), a long short-term memory neural network (LS-MNN) with balanced number of training samples was used by Medvedev et al. achieving more than 90% accuracy ^31^. Zuo et al. ^33^ proposed the CNN-based method for identifying the two kind of HFOs in ripple and fast-ripple separately and achieved average results with sensitivity (77.04% and 83.23% for ripples) and specificity (72.27% and 7 9.36% for fast ripples) compared to four traditional automated methods proposed in the RIPPLELAB toolbox ^32^. The combination of short-time energy (STE) and CNN also used in recent study for identifying HFOs ^62^. In their study, the performance of the system in terms of sensitivity and FDR are used to evaluate their system and compared with three related existing studies ^32, 36, 59^. They achieved higher average results with five adult patients for ripple (Sen: 81.1% and FDR: 30.2%) and fast ripple (Sen: 74.6% and FDR: 6.3%). However, their above studies focused on the detection of HFOs in ripple and fast ripple iEEG data separately and the performance evaluation metrics of their system were mainly used based on their balanced or imbalanced problems. Compared to the above HFOs-related studies, we used only 30-min of iEEG data with SOZ and combined the ripple and fast ripple bands together with the multi-band fashion to identify electrodes related to epileptic events. The average sensitivity, specificity, and Fall-out for individual segment identification with eight patients (including different pathological types with pediatric and adult patients) was 52.70%, 90.75%, and 9.24%, respectively. The average AUC for identifying epileptic focus was 0.86 across eight patients. However, some channels for each patient have created a block of epileptic activity detection (see in Fig. 3). Due to the complex nature of the biological systems, the interictal iEEG are strongly non-stationary, which do not allow the linear methods to adapt perfectly over the whole time windows that is a main reason for creating the block of epileptic activity detection.

In order to achieve a more practical system for real-life applications, we have considered further improvements in the following directions. First, we used only 30-min of signal of the interictal phase, whereas an epileptologist can predict the epileptic focus using the proposed automatic system. We need to expand the automated system using the detection of seizure discharges. Second, the influential parameters to design the system were used based on the previous studies ^30, 53, 63^. The values of the parameters as well as the choice of optimal subbands in multi-band analysis are required to adjust in a data-adaptive nature in order to further improve the system. Third, this study evaluate the subject-dependent system based on the SOZ from the discharges of habitual seizures. Due to subject-specific nature of iEEG signals, the distributions of extracted features among patients were distinct. In machine-learning research, several studies proposed the use of domain transfer learning to adapt the different distributions of features extracted from different subjects ^64, 65^. We strongly believe that domain transfer learning to implement subject-independent system could be one of the best solutions for future study. However, the problem is very challenging due to very different locations of electrodes and subjects-specific nature of epilepsy events. In addition, Islam et al. ^27^ reported that the appropriate selection of operational subbands can significantly improve the system performance due to subject-specific nature of EEG signals. Therefore, the possible extension of this study is to detect the most significant subbands in the high-frequency components, which may further improve our system performance in the future. Thus, there are several avenues for further research to design the automatic system with feature-extraction and classification.

## Conclusion

This study developed an effective epileptic focus detection method from high-frequency components for interictal iEEG data. Eight feature-extraction methods with multi-band analysis were proposed and tested. We evaluated the proposed method for eight epilepsy patients considering different ages and pathological types (adult and pediatric) to investigate efficiency in the high-frequency components (ripple and fast ripple). The detection results were broader around the SOZ electrode ranges for the patient of BOS-FCD type pathology. Moreover, we had the variability of AUC with the patients, in which the pediatric patients have a tendency toward less sensitivity than the adult patients.

## Materials and Methods

### Dataset

More than 100 patients with focal cortical dysplasia (FCD) were studied at the Juntendo University–Epilepsy Center in Tokyo, Japan. This study was approved by the ethics committee of Juntendo University Hospital as well as the Tokyo University of Agriculture and Technology, Japan. All methods were performed in accordance with relevant guidelines and regulations. All the patients signed the informed consent for a research protocol. During pre-surgical evaluation, several non-invasive diagnostic protocols, such as seizure semiological evaluation, interictal scalp EEG, MRI, molecular imaging, and psychomotor-development testing, were performed to determine the electrode locations for each patient. Video-EEG monitoring was also indicated for drug-resistant epilepsy cases as a pre-surgical evaluation.

The subdural electrodes (4-mm diameter and 10-mm distance) (UNIQUE MEDICAL Co, Tokyo, Japan) were implanted and covered almost the entire surface over the FCD and the adjacent cortex. In patients with the bottom of sulcus (BOS) type of dysplasia, the surgeon dissected the cortical sulcus and implanted small electrodes on the vertical sulcus. The iEEGs were acquired using the Neuro Fax digital video EEG system (NIHON-KODEN, Tokyo, Japan) with a sampling rate of 2 kHz. The number of electrodes were defined for each patient based on an epileptologist’s review during iEEG data recording. Among 100 patients, epileptologists selected only eight patients with SOZ and a positive (focal) label was assigned to a channel judged to a seizure onset electrode by epileptologists, and a negative (non-SOZ) label was given to the rest of the channels. Therefore, data on eight patients obtained from SOZ and non-SOZ electrodes were used to evaluate the proposed method. Table 5 shows the summary of the iEEG dataset from the eight patients. The 3D representation of the brain and the electrode positions during recording are shown in Fig. 4 for each patient. The red circle represents the SOZ ^22, 23^ marked by epileptologists.

**Table 5:**
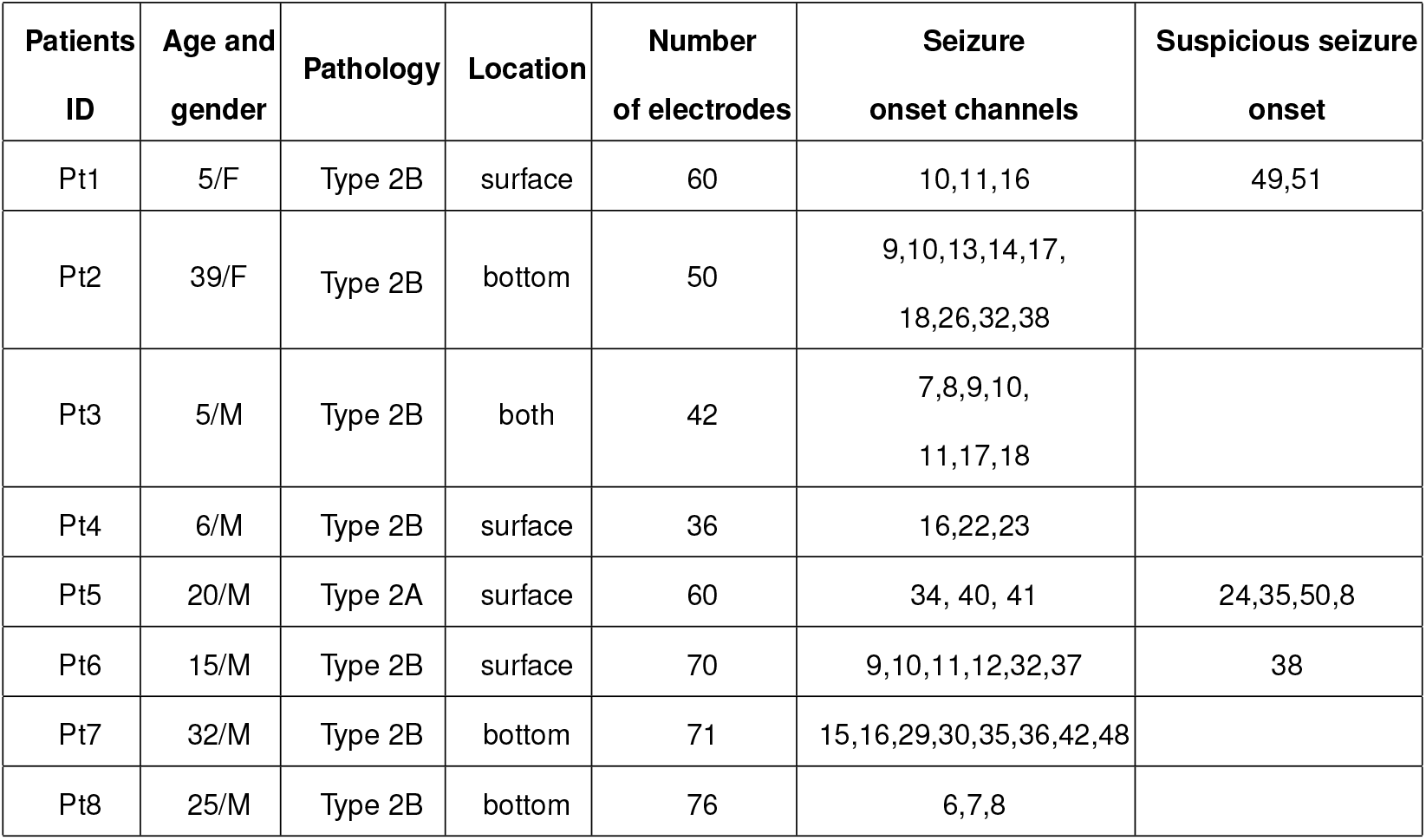
The summary of interictal iEEG data for individual patients with focal cortical dysplasia (FCD). Male and female are indicated as M and F.

**Fig. 4.**
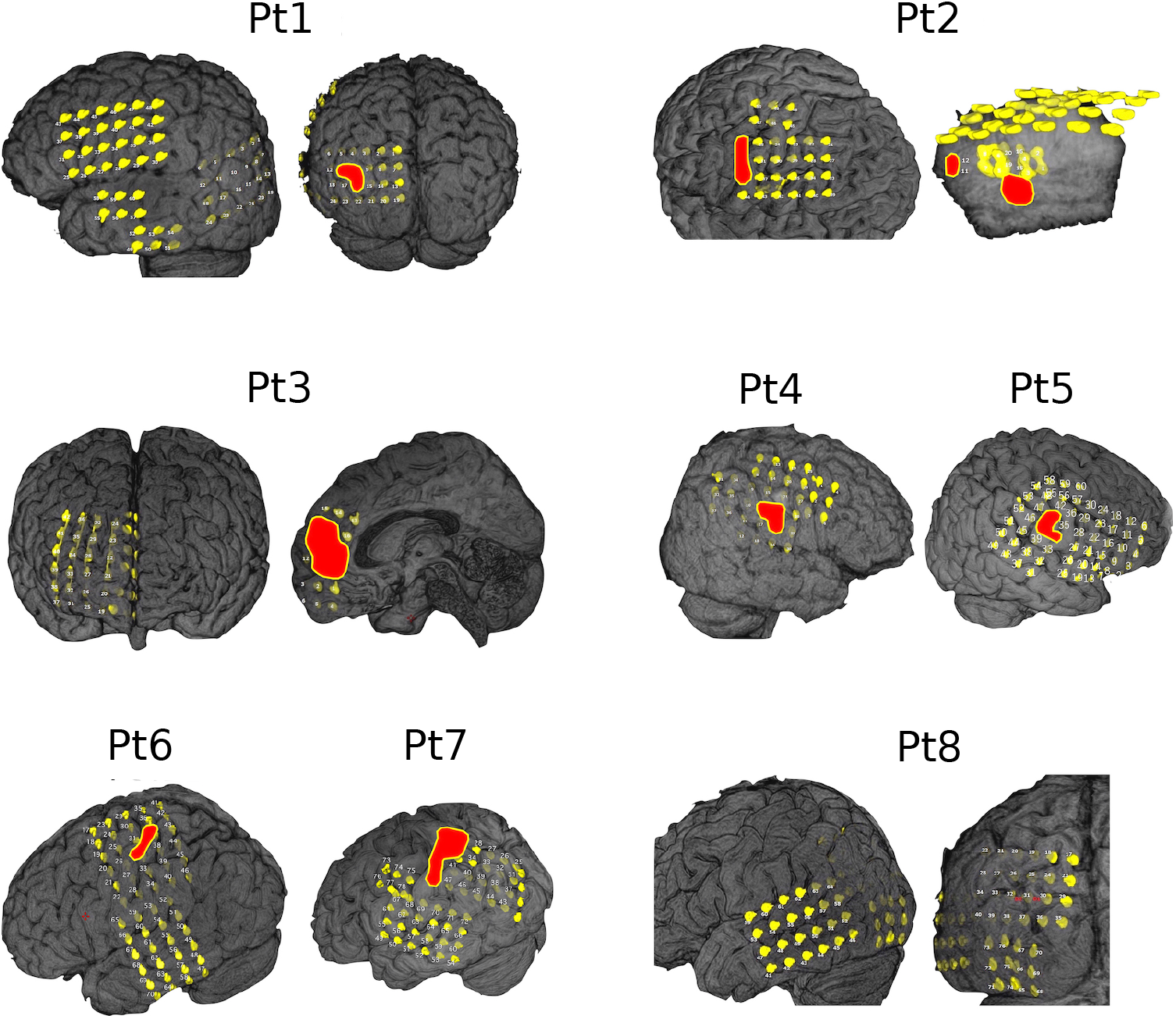
The 3D representation of the brain with interictal electrodes (yellow) of eight patients used from the dataset. The red circle represents the SOZ marked by epileptologists.

### Data Pre-processing

The multi-channel interictal iEEG data were recorded for at least three days until an adequate number of habitual seizures were obtained for analyzing. The epileptologists reviewed the interictal iEEG recordings and annotated the electrodes that had possibly indicated the SOZ. In this study, we used 30-min interictal iEEG data from each patient and split the 30-min iEEG signals into 20-s segments resulting in total 90 segments. The label (SOZ / non-SOZ) was given to each electrode, we assumed that all the segments in SOZ can be considered as focal segments and the segments for other channels were assumed to be non-focal segments. A third-order Butterworth bandpass fitler was applied to extract the high-frequency components (100–600 Hz) from each interictal iEEG segment.

### Multiband Analysis

In practice, the EEG time series that exhibits nonstationary behavior with a variety of neurological events may contain noise that can deteriorate the performance of the system in a single-band approach. Therefore, a filter bank, which is an array of bandpass filters, was applied to decompose an EEG signal into a set of analysis signals exhibiting multiple subband frequency components ^66, 67^. To develop more accurate detection of brain activities related to the specific mental tasks, EEG-based studies proposed the filter-bank method to divide the wide frequency ranges into narrow subbands ^26, 27, 68^. More specific, Higashi et al. proposed a filter-bank approach to improve the performance of MI-BCI, which decomposed the 4–40 Hz frequency ranges into 6 subbands with a bandwidth of 6 Hz each ^69^. Ang et al. divided the similar frequency ranges (4–40 Hz) into narrow subbands with a bandwidth of 4 Hz each ^68^. In signal processing study, the choice of subbands should be as narrow as possible to achieve more accurate detection of automatic system similar to these EEG-BCI ^26, 27, 68, 69^. However, the choice of dividing the wide frequency bands into narrow subbands indeed depends on system performance, real-time applications as well as the reduction of system complexity ^18, 26, 70^. Considering system performance as well as the reduction of the system complexity, the proposed multiband approach divided the high-frequency bands, including ripple (100–250 Hz) and fast ripple (250–600 Hz), into 10 subbands, each of which has a band width of 50 Hz. The subbands are labeled *S*_1_, *S*_2_, …, *S*_*N*_ as shown in Fig. 5, where *N* is the total number of subbands (*N* = 10).

**Fig. 5.**
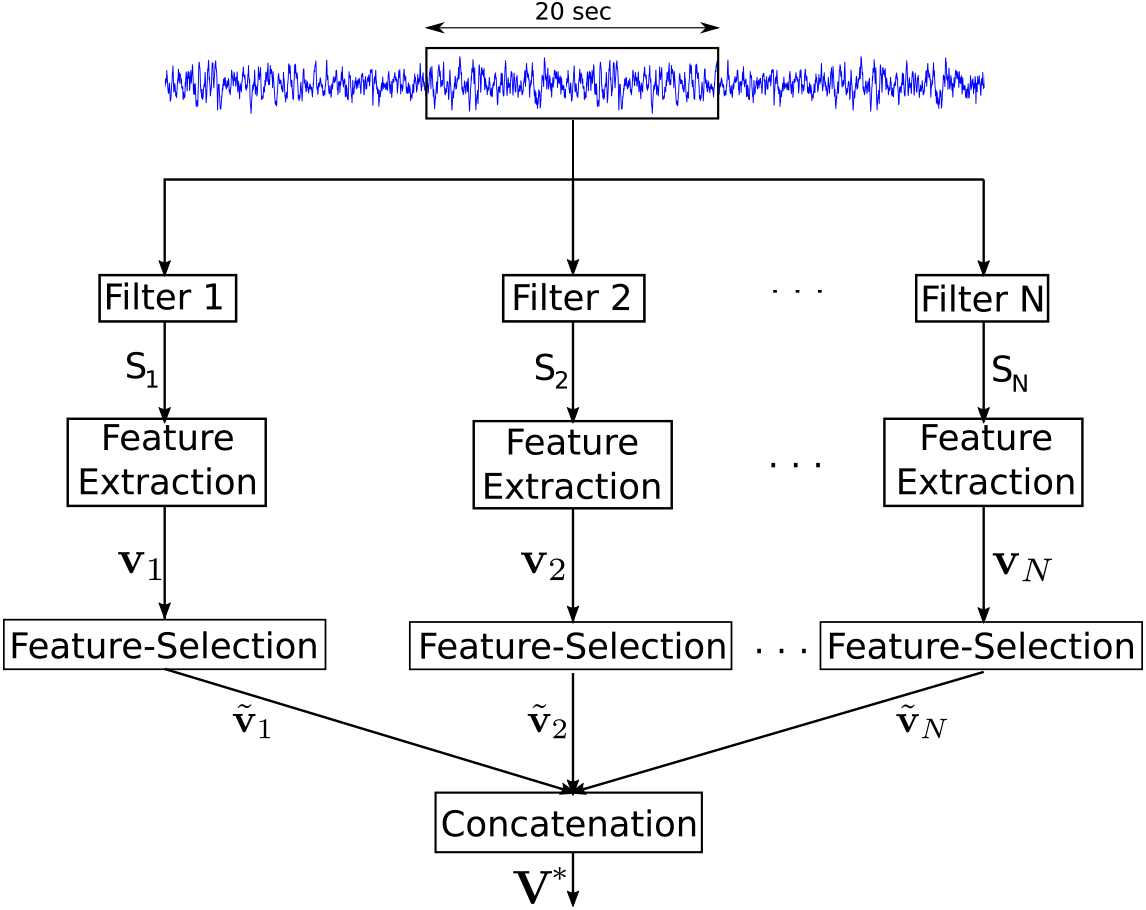
The different components of the proposed system for epileptic focus identification. The values *S*_1_, *S*_2_, …, *S*_*N*_ represent the subbands and *N* represents the total number of subbands.

### Entropy Feature Extraction

To extract features characterizing the complexities of a time-series from interictal iEEG signals, several entropy-based methods are available. It has been reported in different epilepsy studies ^58, 63, 71^ that using a combination of different entropies to extract features from an EEG signal can improve the classification performance. Therefore, the representation of the eight entropies with a multiband approach was chosen to extract features in this study. The details of the eight entropy measures used in this study are summarized in the following sections.

#### Approximate Entropy

Approximate entropy (APE) was first proposed by Pincus et al. to measure the amount of regularity in the time-series ^72^. It is extensively used in many areas of biomedical signal processing, such as EEG ^73^ and ECG signal analysis ^74^. To estimate the approximate entropy (APE) from each segment, let us define as a time series *x*(*i*) of the n-th subbands *S*_*n*_ of each channel. The time series *x*(*i*) can be represented *L* − *d* + 1 vectors as *X*(1), *X*(2), …, *X*(*L* − *d* + 1), where *L* is the length of signal (in our case *L* = 40, 000 for each channel of a segment due to 2 kHz sample frequency). Each *X*(*i*) vector can be expressed as:

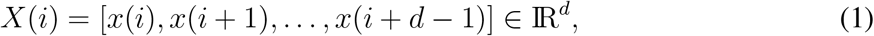

where *d* is the embedding dimension and for each *i*, 1 ≤ *i* ≤ *L* − *d* + 1. APE is defined as:

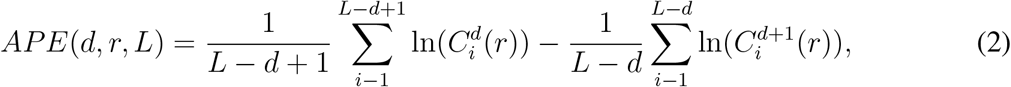

where 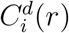 is a correlation integral indicating the probability of the vector *X*(*i*), which remains similar to *X*(*j*) within tolerance limit *r*. The 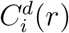 is defined as ^75, 76^:

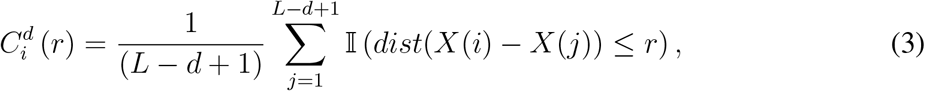

where 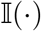 is the indicator function and the *dist*(·) represents the distance between two vectors *X*(*i*) and *X*(*j*). In this study, the value of the *r* parameter is chosen as the 0.2 times the standard deviation of the data, and *d* = 2 used in the study ^72^.

#### Sample Entropy

Sample entropy (Sp) is a modified version of APE intended to resolve a weakness of APE ^77^. The main drawback of APE is a biased estimate due to self-matches of templates. Sp reduces the bias caused by the use of the self matches in the computation of APE ^77^. Sp is defined for a given time series *x*(*i*) as:

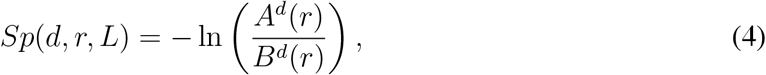

where

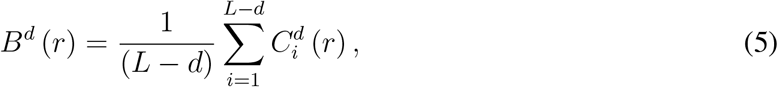

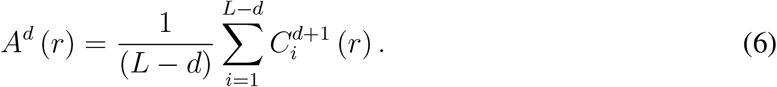

The 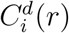 is defined as ^75, 76^:

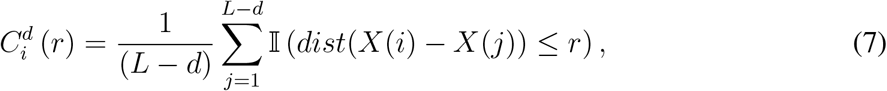

where *X*(*i*) is a vector induced from eq. (1) and 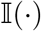 is the indication function to count the true condition number excluding the self-matches 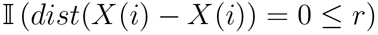 ^72^. In this study, the parameters *r* and *d* were set to the similar to the approximate entropy.

#### Permutation Entropy

Permutation entropy (PE) is a simple and robust method for estimating the complexity of a time series used for automated seizure prediction ^78^. For a given time series *x*(*i*), each vector *X*(*i*) = [*x*(*i*), *x*(*i* + *τ*), …, *x*(*i* + (*d* − 1)*τ*)], where the *d* and *τ* are the embedding dimension and time lag, respectively. Let us define a permutation of [1, 2, …, *d*] by Π = [*j*_1_, *j*_2_, …, *j*_*d*_] in such a way that *x*(*i* + (*j*_1_ − 1)*τ*) ≤ *x*(*i* + (*j*_2_ − 1)*τ*) ≤ … ≤ *x*(*i* + (*j*_*d*_ − 1)*τ*). Then, we can define 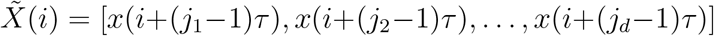. For the set of vectors 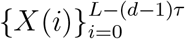, the probability of each possible permutation Π_*k*_ (*k* = 1, 2, …, *d*!) can be introduced as *p*(Π_*k*_) = *C*(Π_*k*_)/(*L* − (*d* − 1)*τ*), where *L* is the length of time series *x*(*i*) and *C*(Π_*k*_) is the number of occurrences of the order pattern Π_*k*_. The PE can be defined as:

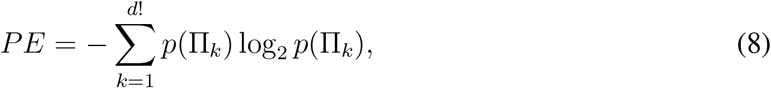

In this study, the parameters *d* and *τ* were set to 3 and 1, respectively.

#### Spectral Entropy

Spectral entropies quantify the complexity of a time series based on the power spectrum ^79^. Several studies have proposed the use of spectral entropy, including Shannon (Sh) and Reny’s entropy (Ren), to characterize the seizure activities ^79–81^. To obtain the power level for each frequency, the Fourier transform of the time series *x*(*i*) is used. The normalization of the power *p*_*f*_ was estimated as:

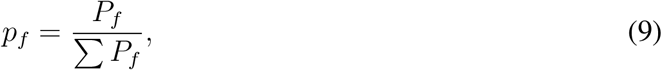

where *P*_*f*_ is the power level of the frequency component. The entropies defined as *Sh* and *Ren* are estimated in follows ^79^:

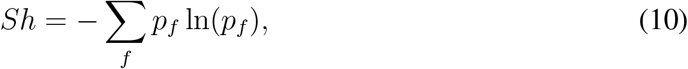

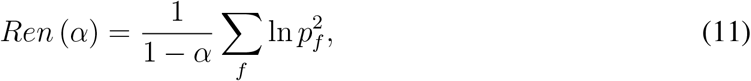

where *α* is the order of Reny’s entropy (*α* = 2).

#### Phase Entropy

Phase entropies are defined through a bispectrum known as higher order spectra ^82^. The bispectrum of a time series *x*(*i*) can be defined as:

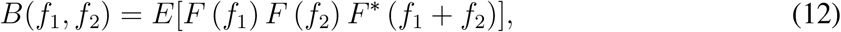

where *E* represents the expectation operator of a random variable. The *F* is the Fourier transform of the time series *x*(*i*) and *F*\* is its conjugate. The two types of phase entropy, *S*_1_ and *S*_2_, can be defined as:

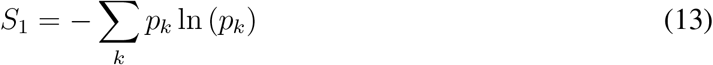

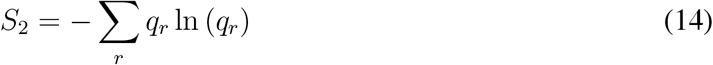

where 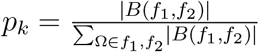 and 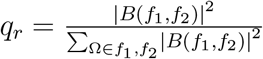.

#### Tsallis Entropy

Tsallis entropy is the generalized version of Shanon entropy and controls the trade off between the contributions from the tails and the main mass of the distribution ^83^. Tsallis entropy is defined as ^83^:

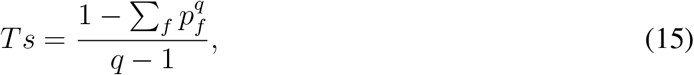

where *p*_*f*_ is the normalization of power computed from the eq. (9) and *q* is a real number, frequently called the entropic-index, that characterizes the degree of non-extensivity of the system^84, 85^. In this study, we set *q* = 2.

### Feature Selection

In machine learning, one of the challenges is the selection of the best feature set from all the available feature space reported in different studies ^27, 86–88^. The selection of entropy features extracted from interictal iEEG data could provide a more accurate classification with respect to the whole set of features. To select more relevant entropy features, sparse LDA is a recently advanced technique ^89, 90^, which reveals discriminant directions of a few variables instead of all the variables used in the standard LDA ^91, 92^. After extracting entropy features from an interictal iEEG segment, the entropies of *n*-th subband for a channel can be defined as:

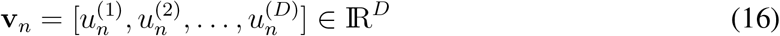

where **v**_*n*_ denotes the combination of entropies (*D* = 8) extracted from the *n*-th subband of a channel using the above feature-extraction methods. We can calculate the entropies for all channels with each segment and finally stacked all of the segments to form the training features *M*_*n*_ ∈ ℝ^*H*×*D*^, where *H* = *ch* × *s* such that *ch* and *s* are the total number of channels and segments, respectively. The sparse LDA criterion from the set of the training features *M*_*n*_ and class *C*_*n*_ for *n*-th subband is defined sequentially as ^93^:

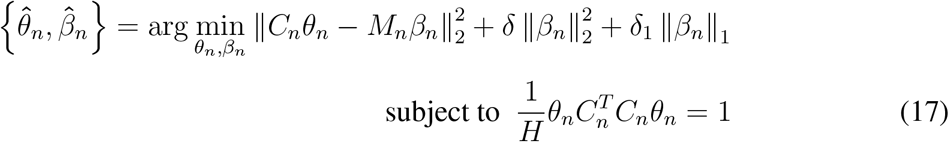

where *θ*_*n*_ = (1, 1)^*T*^ is the initialization vector. The *δ* and *δ*_1_ are tuning parameters used to achieve non-zero elements in each discriminative direction. By solving eq. (17), we will achieve the 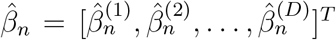. The parameters *δ* and *δ*_1_ were tuned such that 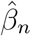 has *G* non-zero elements. Let us define the index of the 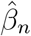 as 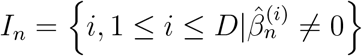. The features 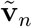 for *n*-th subband with a channel can be defined as:

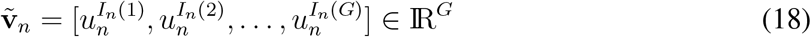

Finally, feature **V**\* is defined by concatenating features 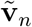 of *N* subbands for a channel as:

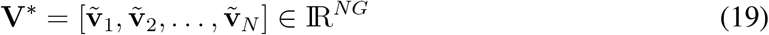

After applying sLDA, the selected training features for *N* subbands can be written as:

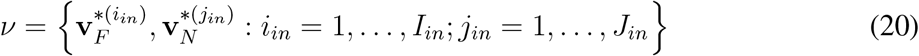

The feature vector of the *i*_*in*_-th sample of SOZ channel is denoted by 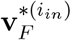 and the feature vector of the *j*_*in*_-th sample of non-SOZ channel is denoted by 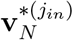. Note that the dataset is generally imbalanced say *I*_*in*_ << *J*_*in*_.

### Imbalanced Learning Problem

In the case of epileptic focus detection, the number of non-SOZ electrodes representing the majority class is much higher than the SOZ electrodes (minority class). This can produce several difficulties in standard machine learning methods due to an imbalance in class distribution and concept complexity ^30, 94, 95^. Therefore, the use of sampling methods in imbalanced learning applications requires the modification of an imbalanced data set by some mechanisms in order to provide a balanced distribution ^94, 95^. Recent studies have shown that a balanced data set with several base classifiers provides improved classification performance compared to an imbalanced data set ^30, 94–96^. In this section, we generate surrogate data using the adaptive synthetic (ADASYN) approach, which is one of the solutions used to solve the imbalanced learning problem. The balance set 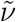 can be defined from the training feature set *ν* induced from eq. (20) as:

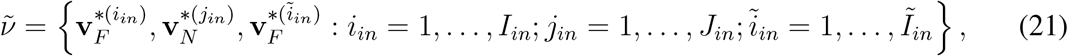

where, 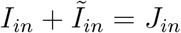. The following algorithm, proposed by He et al. ^30, 94^, is employed here to generate surrogate samples 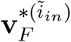.

**Step 1** Calculate the number of synthetic data examples that need to be generated for the entire focal class by:

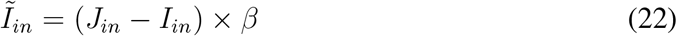

The *β* represents an arbitrary number in the range of 0 to 1 to specify the desired balance level after the synthetic data generation process. We set the *β* in eq. (22) to 1, which corresponds to fully balanced data ^30^.

**Step 2** For each example 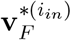 in the focal class, find the *K*-nearest neighbors according to the Euclidean distance and calculate the ratio 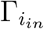 as follows:

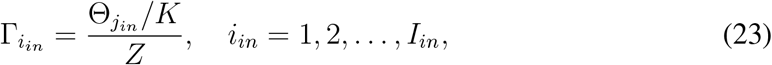

where 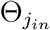 is the number of samples in the *K*-nearest neighbor of 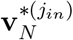 that belong to the non-focal class and *Z* is a normalization factor such that 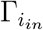 is a distribution function 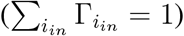.

**Step 3** Determine the number of synthetic samples to be generated for each 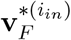 in the focal class as:

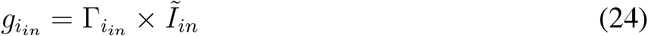

**Step 4** Generate 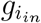 synthetic data samples for each sample of focal class using SMOTE algorithm ^97^ as:

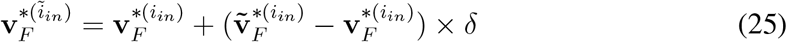

where 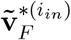 is a randomly chosen focal data example from the *K*-nearest neighbors (*K* = 5) for 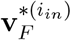 and *δ* denotes the random number belonging to [0,1]. The other parameters for ADASYN were used as default setting ^30^.

### Cross-validation Design

To evaluate the developed system, we needed to divide the data into training and test sets, which was the critical step due to an imbalanced number of focal and non-focal channels. To optimally divide the data into training and test sets, this study proposes k-fold cross-validation technique (*k* = 10) by dividing 90 segments into *k* subsets of equal size. Among the *k* subsets, a single subset is retained for testing the model, and the remaining (*k* − 1) subsets are used as training. As mentioned, ADASYN was applied to highly imbalanced feature sets in the training stage to balance the class features. The cross-validation process is then repeated *k* times and the result of a system is taken by averaging all the runs.

### Performance Measurement for Segments

Instead of using classification accuracy as a system evaluation criterion for imbalanced datasets, a set of assessment metrics related to receiver operating characteristics (ROC) graphs ^29^ were used as performance measurements. Under the imbalanced learning condition, the classification accuracy is not sufficient as a standard performance measure ^29, 98–100^. Therefore, the representation of classification performance can be derived from the confusion matrix, as illustrated in Table 6. Based on Table 6, the evaluation metrics can be defined as:

- Sensitivity (SEN) or recall:

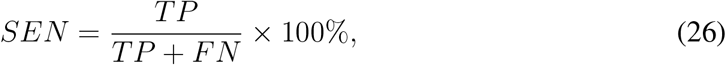

where *TP* is the number of correctly detected segments from the total number of focal segments in the SOZ channels and *FN* indicates the number of incorrectly detected segments from the total number of focal segments in the SOZ channels.
- Specificity (SPE):

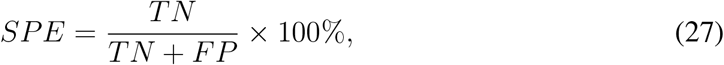

where *TN* is the number of correctly detected segments from the total number of non-focal segments in the non-SOZ channels and *FP* represents the number of incorrectly detected segments from the total number of non-focal segments in the non-SOZ channels.
- Precision or positive predictive value (PPV):

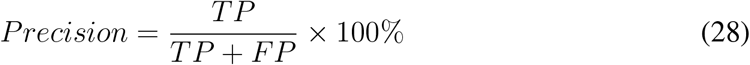
- Fall-out or false positive rate (FPR):

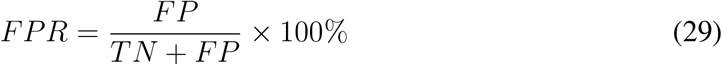
- F_1_ score is the harmonic mean of preision and sensitivity defined as:

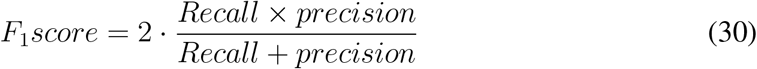

**Table 6:**
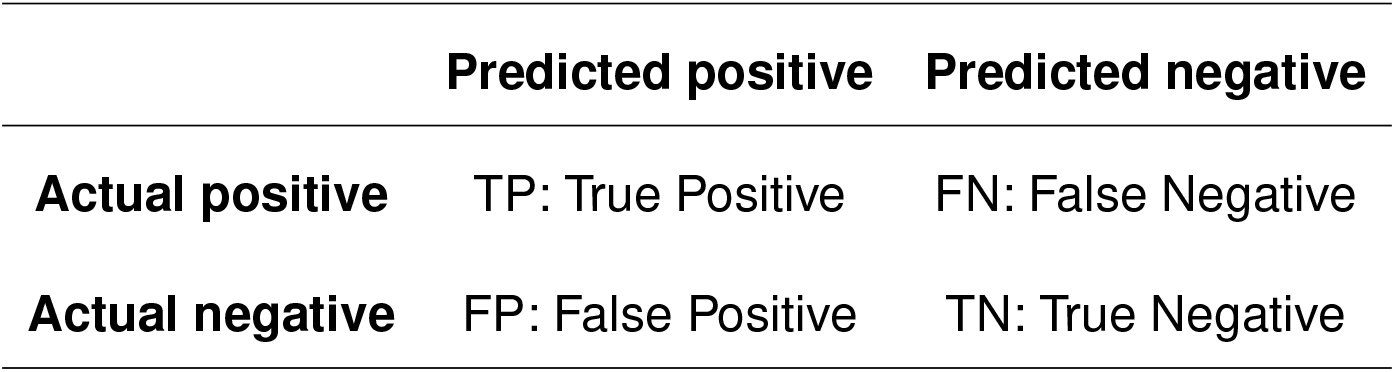
Confusion matrix for a two-class problem.

|  | Predicted positive | Predicted negative |
| --- | --- | --- |
| Actual positive | TP: True Positive | FN: False Negative |
| Actual negative | FP: False Positive | TN: True Negative |

### Performance Measurement for Channels

The performance of each patient to identify epileptic focus was estimated by using AUC-ROC ^101^. The sensitivity (SEN) and false positive rate (FPR) of the channels for each fold were computed using each threshold values (in our case, zero to maximum number of detected focal segments for each fold). After achieving SEN and FPR of the channels with each fold, we estimated the AUC by using trapezoid rule ^101^ and average all the folds to achieve the final results.

## Acknowledgements

We would like to thank Dr. Fabien Lotte (Inria, France) and Dr. Anh Huy Phan (skoltech, Russia) for their discussion and interesting research direction in this work. This work was supported by JST CREST (JPMJMR1784).

## Author Contributions

M.S.A. and K.F. implemented filter-bank and the feature-extraction methods. M.S.A. and M.R.I. conceived the experiments and analysed the results. M.S.A., D.W., M.R.I. and T.T. discussed and implemented the feature selection method. K.F., M.R.I. and T.T. contributed to the technical evaluation. M.S.A., M.R.I. and T.T. wrote the manuscript. A.C. contributed for the validation of the usage of machine learning and signal processing methodologies. H.S. and Y.I. involved to record the interictal iEEG data and contributed to discussion of the clinical evaluation. T.T. supervised the entire project. All authors reviewed the manuscript.

## Competing Interests

The authors declare that they have no competing interests.

## References

1. Fisher, R. S. et al. ILAE official report: A practical clinical definition of epilepsy. Epilepsia 55, 475–482 (2014).

2. Pati, S. & Alexopoulos, A. Pharmacoresistant epilepsy: From pathogenesis to current and emerging therapies. Clevel. Clin. J. Med. 77, 457–467 (2010).

3. Ngugi, A. et al. Incidence of epilepsy. Neurology 77, 1005–1012 (2011).

4. van Mierlo, P. et al. Functional brain connectivity from EEG in epilepsy: Seizure prediction and epileptogenic focus localization. Progress in Neurobiology 121, 19–35 (2014).

5. Yaffe, R. B. et al. Physiology of functional and effective networks in epilepsy. Clinical Neurophysiology 126, 227–236 (2015).

6. Duncan, J. S., Sander, J. W., Sisodiya, S. M. & Walker, M. C. Adult epilepsy. The Lancet 367, 1087–1100 (2007).

7. Miller, J. W. & Shahin, H. Surgical treatment of epilepsy. Epilepsy 19, 730–742 (2013).

8. Worrell, G. A. et al. High-frequency oscillations and seizure generation in neocortical epilepsy. Brain 127, 1496–1506 (2004).

9. Andrzejak, R. G., Schindler, K. & Rummel, C. Nonrandomness, nonlinear dependence, and nonstationarity of electroencephalographic recordings from epilepsy patients. Phys. Rev. E 86, 046206 (2012).

10. Andrzejak, R. G. et al. Indications of nonlinear deterministic and finite-dimensional structures in time series of brain electrical activity: Dependence on recording region and brain state. Phys. Rev. E 64, 061907 (2001).

11. Ortega, G. J., Menendez de la Prida, L., Sola, R. G. & Pastor, J. Synchronization clusters of interictal activity in the lateral temporal cortex of epileptic patients: Intraoperative electrocorticographic analysis. Epilepsia 49, 269–280 (2008).

12. Gajic, D., Djurovic, Z., Gligorijevic, J., Di Gennaro, S. & Savic-Gajic, I. Detection of epileptiform activity in EEG signals based on time-frequency and non-linear analysis. Frontiers in Computational Neuroscience 9, 38 (2015).

13. Lee, S.-H., Lim, J. S., Kim, J.-K., Yang, J. & Lee, Y. Classification of normal and epileptic seizure EEG signals using wavelet transform, phase-space reconstruction, and euclidean distance. Computer Methods and Programs in Biomedicine 116, 10–25 (2014).

14. Srinivasan, V., Eswaran, C. & Sriraam, N. Approximate entropy-based epileptic EEG detection using artificial neural networks. IEEE Transactions on Information Technology in Biomedicine 11, 288–295 (2007).

15. Nicolaou, N. & Georgiou, J. Detection of epileptic electroencephalogram based on permutation entropy and support vector machines. Expert Systems with Applications 39, 202–209 (2012).

16. Song, Y. & Lio, P. A new approach for epileptic seizure detection: sample entropy based feature extraction and extreme learning machine. J. Biomed Sci Eng, 3, 556–567 (2010).

17. Sharma, R. & Pachori, R. B. Classification of epileptic seizures in EEG signals based on phase space representation of intrinsic mode functions. Expert Systems with Applications 42, 1106–1117 (2015).

18. Itakura, T. & Tanaka, T. Epileptic focus localization based on bivariate empirical mode decomposition and entropy. In 2017 Asia-Pacific Signal and Information Processing Association Annual Summit and Conference (APSIPA ASC), 1426–1429 (2017).

19. Andrzejak, R. G., Schindler, K. & Rummel, C. Nonrandomness, nonlinear dependence, and nonstationarity of electroencephalographic recordings from epilepsy patients. Phys. Rev. E 86, 046206 (2012).

20. Bajaj, V. & Pachori, R. B. Epileptic seizure detection based on the instantaneous area of analytic intrinsic mode functions of EEG signals. Biomedical Engineering Letters 3, 17–21 (2013).

21. Zhou, M. et al. Epileptic Seizure Detection Based on EEG Signals and CNN. Frontiers in Neuroinformatics 12, 95 (2018).

22. Jacobs, J. et al. Interictal high-frequency oscillations (80-500 Hz) are an indicator of seizure onset areas independent of spikes in the human epileptic brain. Epilepsia 49, 1893–1907 (2008).

23. Dümpelmann, M., Jacobs, J., Kerber, K. & Schulze-Bonhage, A. Automatic 80-250 Hz ripple high frequency oscillation detection in invasive subdural grid and strip recordings in epilepsy by a radial basis function neural network. Clinical Neurophysiology 123, 1721–1731 (2012).

24. Kerber, K. et al. Differentiation of specific ripple patterns helps to identify epileptogenic areas for surgical procedures. Clinical Neurophysiology 125, 1339–1345 (2014).

25. Ang, K. K., Chin, Z. Y., Wang, C., Guan, C. & Zhang, H. Filter bank common spatial pattern algorithm on BCI competition iv datasets 2a and 2b. Frontiers in Neuroscience 6, 39 (2012).

26. Islam, M. R., Molla, M. K. I., Nakanishi, M. & Tanaka, T. Unsupervised frequency-recognition method of SSVEPs using a filter bank implementation of binary subband cca. Journal of Neural Engineering 14, 026007 (2017).

27. Islam, M. R., Tanaka, T. & Molla, M. K. I. Multiband tangent space mapping and feature selection for classification of EEG during motor imagery. Journal of Neural Engineering 15, 046021 (2018).

28. Zhang, Y., Zhou, G., Jin, J., Wang, X. & Cichocki, A. Optimizing spatial patterns with sparse filter bands for motor-imagery based braincomputer interface. Journal of Neuroscience Methods 255, 85–91 (2015).

29. Fawcett, T. ROC graphs: Notes and practical considerations for researchers. Tech. Rep. (2004).

30. He, H., Bai, Y., Garcia, E. A. & Li, S. Adasyn: Adaptive synthetic sampling approach for imbalanced learning. In 2008 IEEE International Joint Conference on Neural Networks (IEEE World Congress on Computational Intelligence), 1322–1328 (2008).

31. Medvedev, A. V., Agoureeva, G. I. & Murro, A. M. A long short-term memory neural network for the detection of epileptiform spikes and high frequency oscillations. Scientific Reports 9(2019).

32. Navarrete, M., Alvarado-Rojas, C., Michel, L. V. Q. & Valderrama, M. RIPPLELAB: A comprehensive application for the detection, analysis and classification of high frequency oscillations in electroencephalographic signals. PLOS ONE 11, 1–27 (2016).

33. Zuo, R. et al. Automated detection of high-frequency oscillations in epilepsy based on a convolutional neural network. Frontiers in Computational Neuroscience 13, 6 (2019).

34. Edakawa, K. et al. Detection of epileptic seizures using phase-amplitude coupling in intracranial electroencephalography. Scientific Reports 6, 25422 (2016).

35. Cámpora, N. E., Mininni, C. J., Kochen, S. & Lew, S. E. Seizure localization using pre-ictal phase-amplitude coupling in intracranial electroencephalography. Scientific Reports 27(2019).

36. Sato, K. et al. The prehospital predictors of tracheal intubation for in patients who experience convulsive seizures in the emergency department. Intern Med. 16, 21132118 (2017).

37. Pairoj, B., Anannit, V. & Kamornwan, K. Clinical prediction rule of drug resistant epilepsy in children. J Epilepsy Res 5, 84–88 (2015).

38. Anderson, C. F., Moxness, K., Meister, J. & Burritt, M. F. The sensitivity and specificity of nutrition-related variables in relationship to the duration of hospital stay and the rate of complications. Mayo Clinic Proceedings 59, 477–483 (1984).

39. Urrestarazu, E., Jirsch, J. D., LeVan, P. & Hall, J. High-frequency intracerebral EEG activity (100-500 Hz) following interictal spikes. Epilepsia 47, 1465–1476 (2006).

40. Blanco, J. A. et al. Data mining neocortical high-frequency oscillations in epilepsy and controls. Brain 134, 2948–2959 (2011).

41. Zijlmans, M. et al. Ictal and interictal high frequency oscillations in patients with focal epilepsy. Clinical Neurophysiology 122, 664–671 (2011).

42. Jacobs, J. et al. High frequency oscillations (80-500 hz) in the preictal period in patients with focal seizures. Epilepsia 50, 1780–1792 (2009).

43. Urrestarazu, E., Chander, R., Dubeau, F. & Gotman, J. Interictal high-frequency oscillations (100-500 Hz) in the intracerebral EEG of epileptic patients. Brain 130, 2354–2366 (2007).

44. Malinowska, U., Bergey, G. K., Harezlak, J. & Jouny, C. C. Identification of seizure onset zone and preictal state based on characteristics of high frequency oscillations. Clinical Neurophysiology 126, 1505–1513 (2015).

45. Jacobs, J. et al. Interictal high-frequency oscillations (80-500 hz) are an indicator of seizure onset areas independent of spikes in the human epileptic brain. Epilepsia 49, 1893–1907 (2008).

46. Staba, R. J., Wilson, C. L., Bragin, A., Fried, I. & Engel, J. Quantitative analysis of high-frequency oscillations (80-500 Hz) recorded in human epileptic hippocampus and entorhinal cortex. Journal of Neurophysiology 88, 1743–1752 (2002).

47. Gardner, A. B., Worrell, G. A., Marsh, E., Dlugos, D. & Litt, B. Human and automated detection of high-frequency oscillations in clinical intracranial EEG recordings. Clinical Neurophysiology 118, 1134–1143 (2007).

48. Jiang, C. et al. Determining the quantitative threshold of high-frequency oscillation distribution to delineate the epileptogenic zone by automated detection. Frontiers in Neurology 9, 889 (2018).

49. Liu, S. et al. Stereotyped high-frequency oscillations discriminate seizure onset zones and critical functional cortex in focal epilepsy. Brain 141, 713–730 (2018).

50. Quitadamo, L. R. et al. EPINETLAB: A software for seizure-onset zone identification from intracranial EEG signal in epilepsy. Front Neuroinformat 12(2018).

51. Guo, L., Rivero, D. & Pazos, A. Pileptic seizure detection using multiwavelet transform based approximate entropy and artificial neural networks. Journal of Neuroscience Methods 193, 156–163 (2010).

52. Wang, D., Miao, D. & Xie, C. Best basis-based wavelet packet entropy feature extraction and hierarchical EEG classification for epileptic detection. Expert Systems with Applications 38, 14314–14320 (2011).

53. Sharma, R., Pachori, R. B. & Acharya, U. R. An integrated index for the identification of focal electroencephalogram signals using discrete wavelet transform and entropy measures. Entropy 17, 5218–5240 (2015).

54. Mursalin, M., Zhang, Y., Chen, Y. & Chawla, N. V. Automated epileptic seizure detection using improved correlation-based feature selection with random forest classifier. Neurocomputing 241, 204–214 (2017).

55. Shoeb, A. & Guttag, J. Application of machine learning to epileptic seizure detection. In Proceedings of the 27th International Conference on International Conference on Machine Learning, ICML’10, 975–982 (Omnipress, USA, 2010).

56. Ullah, I., Hussain, M., ul Haq Qazi, E. & Aboalsamh, H. An automated system for epilepsy detection using EEG brain signals based on deep learning approach. Expert Systems with Applications 107, 61–71 (2018).

57. Acharya, U. R., Oh, S. L., Hagiwara, Y., Tan, J. H. & Adeli, H. Deep convolutional neural network for the automated detection and diagnosis of seizure using EEG signals. Computers in Biology and Medicine 100, 270–278 (2018).

58. Itakura, T. & Tanaka, T. Epileptic focus localization based on bivariate empirical mode decomposition and entropy. In 2017 Asia-Pacific Signal and Information Processing Association Annual Summit and Conference (APSIPA ASC), 1426–1429 (2017).

59. Jrad, N. et al. Automatic detection and classification of high-frequency oscillations in depth-EEG signals. IEEE Transactions on Biomedical Engineering 64, 2230–2240 (2017).

60. Guo, J. et al. A stacked sparse autoencoder-based detector for automatic identification of neuromagnetic high frequency oscillations in epilepsy. IEEE Transactions on Medical Imaging 37, 2474–2482 (2018).

61. Johansen, A. R. et al. Epileptiform spike detection via convolutional neural networks. In 2016 IEEE International Conference on Acoustics, Speech and Signal Processing (ICASSP), 754–758 (2016).

62. Lai, D. et al. Automated detection of high frequency oscillations in intracranial EEG using the combination of short-time energy and convolutional neural networks. IEEE Access 7, 82501–82511 (2019).

63. Sharma, R., Pachori, R. B. & Acharya, U. R. Application of entropy measures on intrinsic mode functions for the automated identification of focal electroencephalogram signals. Entropy 17, 669–691 (2015).

64. Pan, S. J. & Yang, Q. A survey on transfer learning. IEEE Transactions on Knowledge and Data Engineering 22, 1345–1359 (2010).

65. Lotte, F. & Guan, C. Learning from other subjects helps reducing brain-computer interface calibration time. In 2010 IEEE International Conference on Acoustics, Speech and Signal Processing, 614–617 (2010).

66. Vetterli, M. & Herley, C. Wavelets and filter banks: theory and design. IEEE Transactions on Signal Processing 40, 2207–2232 (1992).

67. Boashash, B. (ed.) Time-frequency signal analysis and processing (Academic Press, Oxford, 2016).

68. Ang, K. K., Chin, Z. Y., Wang, C., Guan, C. & Zhang, H. Filter bank common spatial pattern algorithm on BCI competition IV datasets 2a and 2b. Frontiers in Neuroscience 6, 39 (2012).

69. Higashi, H. & Tanaka, T. Simultaneous design of fir filter banks and spatial patterns for eeg signal classification. IEEE Transactions on Biomedical Engineering 60, 1100–1110 (2013).

70. Chen, X. et al. High-speed spelling with a noninvasive brain–computer interface. Proceedings of the National Academy of Sciences 112, E6058–E6067 (2015).

71. Sharma, R. & Pachori, R. B. Classification of epileptic seizures in EEG signals based on phase space representation of intrinsic mode functions. Expert Systems with Applications 42, 1106–1117 (2015).

72. Pincus, S. Approximate entropy (apen) as a complexity measure. Chaos: An Interdisciplinary Journal of Nonlinear Science 5, 110–117 (1995).

73. Abasolo, D. et al. Analysis of regularity in the EEG background activity of Alzheimer’s disease patients with approximate entropy. Clinical Neurophysiology 116, 1826–1834 (2005).

74. Acharya, U. R. et al. Linear and nonlinear analysis of normal and cad-affected heart rate signals. Computer Methods and Programs in Biomedicine 113, 55–68 (2014).

75. Richman, J. S. & Moorman, J. R. Physiological time-series analysis using approximate entropy and sample entropy. American Journal of Physiology-Heart and Circulatory Physiology 278, H2039–H2049 (2000).

76. Cui, D. et al. Analysis of entropies based on empirical mode decomposition in amnesic mild cognitive impairment of diabetes mellitus. Journal of Innovative Optical Health Sciences 08, 1550010 (2015).

77. Sokunbi, M. O. et al. Nonlinear complexity analysis of brain fMRI signals in schizophrenia. PLOS ONE 9, 1–10 (2014).

78. Bandt, C. & Pompe, B. Permutation entropy: A natural complexity measure for time series. Phys. Rev. Lett. 88, 174102 (2002).

79. Vanluchene, M. D. et al. Spectral entropy as an electroencephalographic measure of anesthetic drug effecta comparison with bispectral index and processed midlatency auditory evoked response. Anesthesiology 101, 34–42 (2004).

80. Blanco, S., Garay, A. & Coyulombie, D. Comparison of frequency bands using spectral entropy for epileptic seizure prediction. ISRN Neurology 2013, 287–327 (2013).

81. Mirzaei, A., Ayatollahi, A., Gifani, P. & Salehi, L. EEG analysis based on wavelet-spectral entropy for epileptic seizures detection. In 2010 3rd International Conference on Biomedical Engineering and Informatics, vol. 2, 878–882 (2010).

82. Nikias, C. L. & Mendel, J. M. Signal processing with higher-order spectra. IEEE Signal Processing Magazine 10, 10–37 (1993).

83. Tsallis, C. Computational applications of nonextensive statistical mechanics. Journal of Computational and Applied Mathematics 227, 51–58 (2009). Special Issue of Proceedings of NUMAN 2007 Conference: Recent Approaches to Numerical Analysis: Theory, Methods and Applications.

84. Tsallis, C. Computational applications of nonextensive statistical mechanics. J. Comput. Appl. Math. 227, 51–58 (2009).

85. Shannon, C. E. A mathematical theory of communication. SIGMOBILE Mob. Comput. Commun. Rev. 5, 3–55 (2001).

86. Chen, Y. M., Lin, P., He, J. Q. & Li, X. L. Combination of the manifold dimensionality reduction methods with least squares support vector machines for classifying the species of sorghum seeds. Scientific Reports 6(2016).

87. John, P. C. & Byron, M. Y. Dimensionality reduction for large-scale neural recordings. Nature neuroscience 17, 1500–1509 (2014).

88. Uehara, T., Sartori, M., Tanaka, T. & Fiori, S. Robust averaging of covariances for eeg recordings classification in motor imagery brain-computer interfaces. Neural Computation 29, 1631–1666 (2017). PMID: 28410052.

89. Clemmensen, L., Hastie, T., Witten, D. & Ersbll, B. Sparse discriminant analysis. Technometrics 53, 406–413 (2011).

90. Zhang, Y. et al. Aggregation of sparse linear discriminant analysis for event-related potential classification in brain-computer interface. International Journal of Neural Systems 24, 1450003 (2014).

91. Lei, Z., Liao, S. & Li, S. Z. Efficient feature selection for linear discriminant analysis and its application to face recognition. In Proceedings of the 21st International Conference on Pattern Recognition (ICPR2012), 1136–1139 (2012).

92. Li, S. Z., Chu, R., Liao, S. & Zhang, L. Illumination invariant face recognition using near-infrared images. IEEE Transactions on Pattern Analysis and Machine Intelligence 29, 627–639 (2007).

93. Sjöstrand, K., Clemmensen, L. H., Larsen, R., Einarsson, G. & Ersbøll, B. SpaSM: A MATLAB toolbox for sparse statistical modeling. Journal of Statistical Software 84, 1–24 (2018).

94. He, H. & Garcia, E. A. Learning from imbalanced data. IEEE Transactions on Knowledge and Data Engineering 21, 1263–1284 (2009).

95. Wang, S., Minku, L. L. & Yao, X. A systematic study of online class imbalance learning with concept drift. IEEE Transactions on Neural Networks and Learning Systems 29, 4802–4821 (2018).

96. Santoso, B., Wijayanto, H., Notodiputro, K. A. & Sartono, B. Synthetic over sampling methods for handling class imbalanced problems : A review. IOP Conference Series: Earth and Environmental Science 58, 012031 (2017).

97. Chawla, N., Bowyer, K., Hall, L. & Kegelmeyer, W. SMOTE: Synthetic minority over-sampling technique. J. Artificial Intelligence Research 16, 321–357 (2002).

98. Guo, H. & Viktor, H. L. Learning from imbalanced data sets with boosting and data generation: The DataBoost-IM approach. ACM SIGKDD Explor. Newsl. 6, 30–39 (2004).

99. Kubat, M., Holte, R. C. & Matwin, S. Machine learning for the detection of oil spills in satellite radar images. Machine Learning 30, 195–215 (1998).

100. Kubat, M. & Matwin, S. Addressing the curse of imbalanced training sets: One-sided selection. In Proceedings of the Fourteenth International Conference on Machine Learning, 179–186 (Morgan Kaufmann, 1997).

101. Fawcett, T. An introduction to ROC analysis. Pattern Recognition Letters 27, 861–874 (2006).

102. Lüders, H. O., Najm, I., Nair, D., Widdess-Walsh, P. & Bingman, W. The epileptogenic zone: General principles. Epileptic Disord. 8, 1–9 (2006).

